# Rapid discovery of antiviral targets through dimensionality reduction of genome-scale metabolic models

**DOI:** 10.1101/2025.10.02.679942

**Authors:** Yong-ki Lee, Seongmo Kang, JinA Lim, Kanghee Kim, Se-Mi Kim, Mark Anthony B. Casel, Issac Choi, Young Ki Choi, Hyun Uk Kim, Yoosik Kim

## Abstract

The COVID-19 pandemic underscored the urgent need for rapid and broadly applicable strategies to identify antiviral targets against emerging pathogens. Conventional approaches, which rely on detailed viral characterization and large-scale drug screening, are too slow to address this challenge. Here, we introduce a transcriptome-based computational framework that integrates genome-scale metabolic models with dimensionality reduction to uncover host metabolic vulnerabilities that support viral replication. Applying this approach to bulk and single-cell RNA-seq data from HCoV-OC43–infected cells and organoids identified oxidative phosphorylation as a key vulnerability, and pharmacological inhibition of complex I effectively curtailed viral replication. Extending the framework to SARS-CoV-2 and MERS-CoV revealed pyrimidine catabolism as a conserved antiviral pathway, with inhibition of its rate-limiting enzyme DPYD suppressing replication in organoid models. Re-analysis of SARS-CoV-2 patient metabolome data further confirmed elevated DPYD activity, underscoring its clinical relevance. Together, these findings establish a generalizable and rapid strategy for host-directed antiviral discovery, providing a foundation for precision therapeutics and pandemic preparedness.

**Significance Statement:** Host-directed antiviral therapies offer several advantages in antiviral research, but identifying key host factors poses a significant challenge. By integrating genome-scale metabolic models with single-gene knockout simulation and dimensionality reduction, we developed a computational framework based on single and bulk RNA-seq data that can systematically pinpoint host pathways whose downregulation is predicted to rewire virus-induced metabolic alterations. Applying this approach to multiple human coronaviruses reveals unique metabolic vulnerabilities, and we experimentally demonstrate that inhibiting these host metabolic pathways reduces viral replication. This framework provides a generalizable antiviral strategy to discern effective targets and can be further extended to investigate virus–host metabolic interactions.

## Introduction

The COVID-19 pandemic has sparked broad interest in developing antiviral strategies to combat emerging and recurring viral threats. Extensive research into the replication mechanisms of SARS-CoV-2 has facilitated both the discovery of novel antiviral drugs and repurposing efforts, exemplified by Nirmatrelvir and Molnupiravir, respectively. In conjunction with mRNA vaccines, these therapeutic interventions have played a pivotal role in shifting the pandemic to its current endemic phase. However, this transition has come at the cost of substantial human loss and profound social and economic burden, highlighting the need for more efficient and prompt response systems. In this context, antiviral research is expanding into underexplored perspectives, one of which focuses on modifying host metabolism to hinder viral replication.

In infected cells, viruses alter host cells’ metabolism to facilitate their reproduction (1, 2). Glucose and fatty acid metabolism have been identified as critical regulatory pathways to support replication of numerous viruses (3), and a broad range of host metabolic processes have been reported to be altered in response to viral infection (4, 5). Inspired by these findings, several studies have modulated specific metabolic pathways, including pyrimidine (6, 7), glutamine (8), lipid (9), and fatty acid (10, 11) synthesis, to inhibit viral replication. Other studies have focused on individual metabolites, showing that specific molecules can influence viral infectivity and pathogenesis (12, 13). However, identifying metabolic targets requires iterative screening and mechanistic insight into virus–host interactions, which poses challenges for rapid responses to emerging viruses.

To address this limitation, *in silico* approaches, such as virtual screening, have been explored (14, 15). Of these, the genome-scale metabolic model (GEM) stands out as a powerful tool for deciphering complex metabolic networks of organisms through genome-protein-reaction (GPR) associations (16). Multi-omics data from various biological samples can be integrated to generate context-specific GEMs (17), which can be used to analyze metabolic flux profiles and predict the effects of single-gene inhibition on overall metabolic fluxes (18). Furthermore, these models can be used to identify biomarkers or discover therapeutic targets (19–23). In the context of viral infection, GEMs have been employed to characterize infection-triggered metabolic alterations and to identify host enzymes that affect viral replication (24–28). Nevertheless, most studies have primarily focused on reducing total viral biomass or on specific metabolic pathways with increased flux in infected cells, with less emphasis on global changes in the host metabolism induced by viruses.

Here, we introduce a computational framework that integrates GEMs with Uniform Manifold Approximation and Projection (UMAP) to capture the metabolic state of virus-infected cells and discover host enzymes whose inhibition is predicted to revert the infected metabolic profile toward that of uninfected cells (Fig. 1). We first validated this approach using publicly available RNA-seq datasets from HCoV-OC43– and HCoV-229E–infected cells, confirming its ability to identify relevant metabolic perturbations. We next extended the analysis to single-cell RNA-seq (scRNA-seq) profiles generated from human bronchial epithelial (HBE) organoids infected with HCoV-OC43, demonstrating its applicability in complex tissue models. Finally, we applied the framework to two high-priority coronaviruses lacking effective treatments, SARS-CoV-2 and MERS-CoV, uncovering conserved metabolic vulnerabilities. Collectively, this strategy provides a generalizable, transcriptome-based platform for dissecting virus–host metabolic interactions and rapidly identifying antiviral targets without the need for prior viral characterization or chemical screening.

**Fig. 1.**
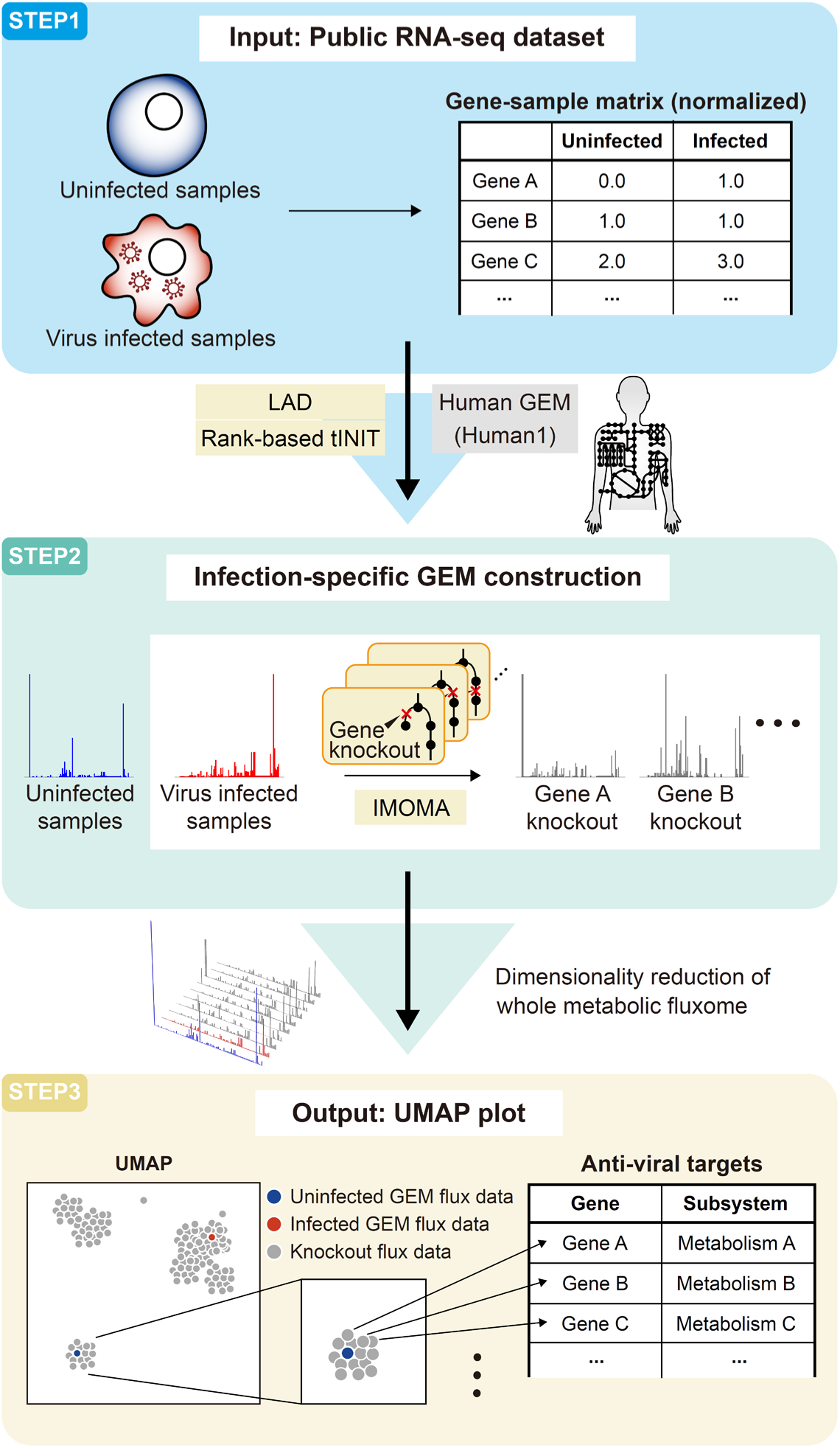
Schematic workflow of the computational framework for antiviral target identification. A schematic workflow illustrating the framework for identifying antiviral targets using transcriptomics datasets. The framework consists of three key steps. First, a normalized gene-sample matrix derived from an RNA-seq dataset containing both uninfected and virus-infected samples is used as the input. Human-GEM (Human1) serves as the template, and infection-specific GEMs are constructed using the rank-based tINIT. Flux profiles (fluxomes) are inferred by the LAD. Second, single-gene knockout simulations are performed on the virus-infected GEMs, and flux changes after gene knockout are predicted using the lMOMA method. Third, all fluxomes (uninfected, infected, and knockouts) are embedded via UMAP to identify genes whose knockout fluxomes co-cluster with that of the uninfected sample. These genes are considered potential antiviral targets.

## Results

### Antiviral Target Discovery via Metabolic Alterations

We first utilized publicly available RNA-seq data to demonstrate the feasibility of the proposed approach for discovering antiviral targets through metabolic alteration. We generated infection-specific GEMs using datasets from two cell lines infected with common human coronaviruses (HCoVs) (MRC-5 lung fibroblast cells infected with HCoV-229E and A549-ACE2 lung adenocarcinoma cells infected with HCoV-OC43) (Fig. 2*A*). For comparison, we also generated a “control GEM” reconstructed from pre-infection (uninfected) data. Moreover, the infected GEM is further utilized as a basis for single-gene knockout simulations discussed below. We then inferred the metabolic flux profiles for each GEM using the least absolute deviation (LAD) (29) and subsequently employed UMAP for dimensionality reduction to simplify the complexity of the flux profiles of all the GEMs. The UMAP visualization represents an individual sample as a separate point, with each point encompassing the entire metabolic flux profile of the sample. We found that both RNA-seq data exhibited distinct clustering between the flux profiles of control and infected cells (Fig. 2*B*, highlighted in blue and red boxes). These results indicate that viral infection causes significant metabolic changes in host cells, enabling the identification of potential targets to revert the cells to their uninfected state.

**Fig. 2.**
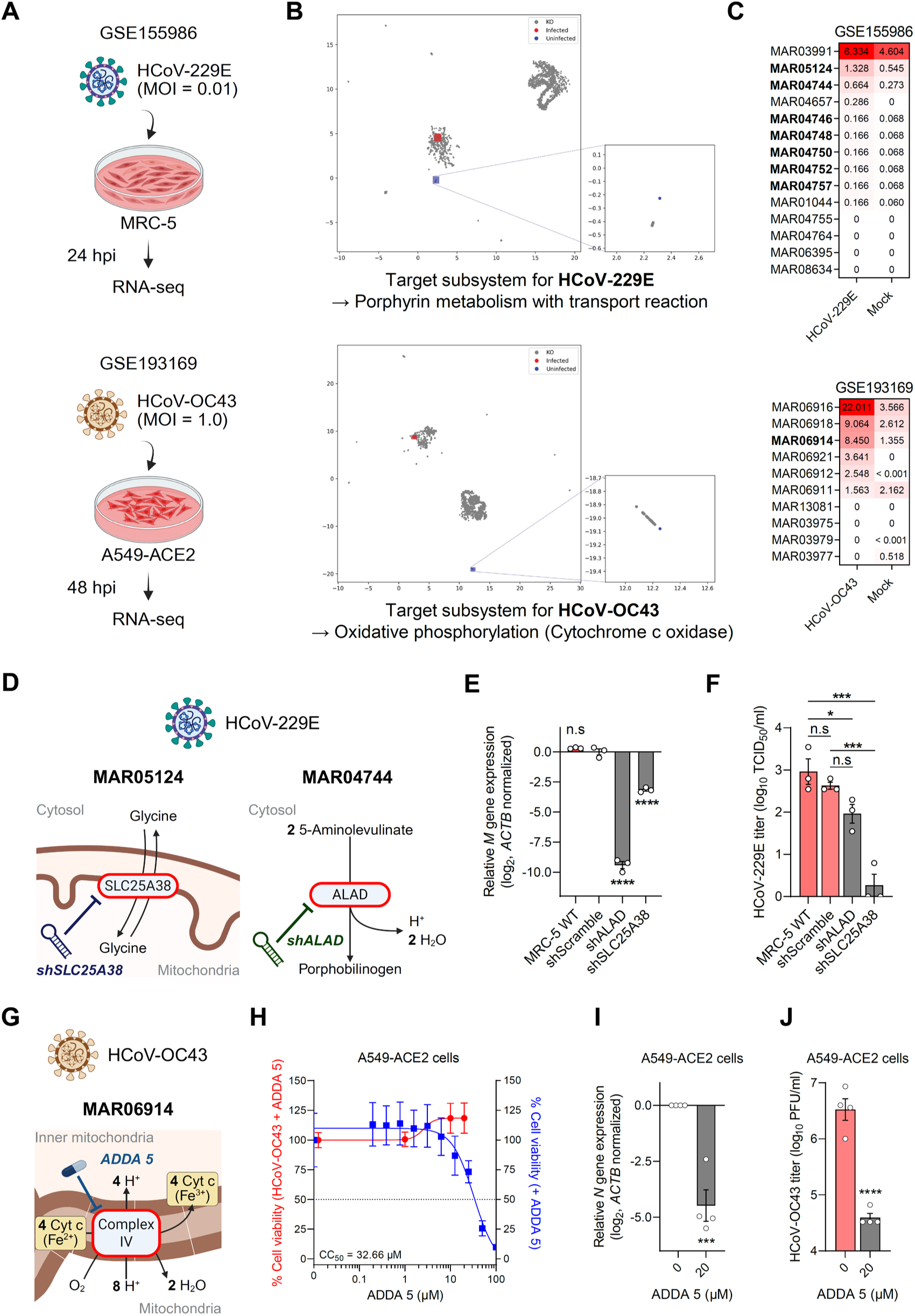
Discovery of antiviral targets for HCoV-OC43 and HCoV-229E. **(A)** Graphical summary of the experimental design for two public RNA-seq datasets, GSE155986 and GSE193169, used as input data for the computational framework to identify antiviral targets. Images were created with BioRender.com. **(B)** UMAP plots showing the fluxome comparisons across all GEMs generated using the GSE155986 or GSE193169 dataset. The uninfected sample is represented in blue, the virus-infected sample in red, and the single-gene knockout simulation results as gray dots. Detailed gene lists are provided in Table S1. **(C)** Metabolic flux profiles of the porphyrin subsystem in HCoV-229E–infected samples and the OXPHOS subsystem in HCoV-OC43–infected samples, compared with uninfected controls. Predicted antiviral target reactions are highlighted in bold. **(D)** Schematic representation of reactions involving SLC25A38 and ALAD with shRNA-mediated inhibition indicated (adapted from https://metabolicatlas.org/). Images were created with BioRender.com. **(E)** Intracellular HCoV-229E membrane (*M*) gene expression at 24 hpi was quantified via RT-qPCR (n = 3). *ACTB* was used as the normalization control. Statistical significance was calculated using one-way ANOVA with Dunnett’s multiple comparison test. **(F)** Extracellular HCoV-229E titers at 24 hpi were measured by the endpoint dilution assay (n = 3). Statistical significance was calculated using one-way ANOVA with Dunnett’s multiple comparison test. TCID_50_: tissue culture infectious dose 50%. **(G)** Schematic representation of the OXPHOS complex IV (cytochrome c oxidase) reaction and its inhibition by ADDA 5 indicated (adapted from https://metabolicatlas.org/). Images were created with BioRender.com. Cyt c: cytochrome c. **(H)** Dose–response curves for ADDA 5 in A549-ACE2 cells, showing viability of uninfected cells (blue, n = 4) and HCoV-OC43–infected cells (red, n = 2). Cell viability was measured after 48 h of treatment. Curves were fitted by nonlinear four-parameter sigmoidal regression in GraphPad Prism. **(I)** Intracellular HCoV-OC43 nucleocapsid (*N*) gene expression at 48 hpi was quantified via RT-qPCR (n = 4). *ACTB* was used as the normalization control. Statistical significance was calculated using a one-tailed Student’s t-test. **(J)** Extracellular HCoV-OC43 titers at 48 hpi were measured by the plaque assay (n = 4). Statistical significance was calculated using a one-tailed Student’s t-test. PFU: plaque-forming unit.

We then performed single-gene knockout simulations on infected GEMs, followed by UMAP for dimensionality reduction, to investigate the metabolic flux differences induced by gene perturbations. Our analysis revealed several single-gene knockout flux profiles that were clustered with that of the uninfected control GEM (Fig. 2*B*, magnified blue boxes). We considered these genes as potential antiviral targets, as their deletion is predicted to shift the overall metabolic flux profiles of infected cells toward a state resembling that of uninfected cells. Indeed, we have previously experimentally validated that this approach can successfully impact overall metabolism and influence the cellular response to chemotherapy drugs in breast cancer cells, serving as a model system (30).

A detailed examination of these profiles revealed that genes in this cluster were functionally related. The subsystem of target genes in HCoV-229E was predominantly associated with porphyrin metabolism (*SI Appendix*, Table S1), while those in HCoV-OC43 were related to oxidative phosphorylation (OXPHOS) (*SI Appendix*, Table S2). A comparison of GEMs between uninfected and infected cells confirmed that the target reaction flux was elevated following viral infection (Fig. 2*C*). Of note, increased flux was not the primary criterion for target selection in our framework. Yet, many of the candidate genes exhibited increased flux upon infection, suggesting that they may play an essential role in metabolic alterations during infection. Interestingly, differentially expressed gene (DEG), gene ontology (GO) and gene set enrichment analysis (GSEA) of the RNA-seq data did not show porphyrin metabolism and oxidative phosphorylation as significantly upregulated pathways (*SI Appendix*, Fig. S1*A*–*D*), indicating that our approach suggests potential regulators of infected cells’ metabolism that cannot be easily found by the conventional analysis of differential gene expression from RNA-seq data.

Based on these findings, we proceeded to investigate the potential impact of metabolic pathways predicted from the framework on viral infection. Porphyrin is a heterocyclic ring compound composed of four pyrrole subunits connected by methine bridges, serving as a framework for heme synthesis. Since no small molecule inhibitors targeting porphyrin metabolism have been discovered to date, short-hairpin RNA (shRNA) was utilized as an alternative approach. We selected aminolevulinate dehydratase (*ALAD*) and solute carrier family 25 member 38 (*SLC25A38*), which are involved in the initial process of porphyrin biosynthesis (31), as knockdown targets among the seven candidate genes (Fig. 2*D*). To mimic a single-gene knockout simulation, MRC-5 cells were transduced with lentiviral vectors encoding shALAD or shSLC25A38 RNAs (hereafter referred to as shALAD and shSLC25A38 cells, respectively). Comparison of their gene expression levels with those of shScramble cells confirmed efficient knockdown (*SI Appendix*, Fig. S2*A*). The impact of target suppression on HCoV-229E replication was evaluated by infecting shScramble, shALAD, and shSLC25A38 cells with HCoV-229E under the same conditions as dataset GSE155986. To account for indirect shRNA-related effects on viral replication, we analyzed viral replication in MRC-5 cells with or without shRNA transduction. A comparative analysis of intracellular viral gene levels revealed a significant reduction in viral gene expression in shALAD and shSLC25A38 cells, whereas no substantial differences were detected between shScramble and no transduction cells (Fig. 2*E*). The measurement of extracellular viral load from cell supernatants also showed no significant difference between shScramble and no transduction samples. In contrast, a decreased viral titer was observed in shALAD and shSLC25A38 cells, with a more pronounced log reduction (2.33) in the shSLC25A38 sample (Fig. 2*F*).

The anti-HCoV-OC43 target genes identified by our computational framework were strongly associated with OXPHOS, specifically the subunits of complex IV (cytochrome c oxidase, CIV). CIV transfers electrons from cytochrome c to oxygen, generating the proton gradient essential for ATP synthesis (Fig. 2*G*). We found that ADDA 5 and chlorpromazine are highly selective inhibitors of CIV (32, 33). Although chlorpromazine is an FDA-approved psychotropic drug, its antiviral properties have already been demonstrated (34, 35) through the inhibition of clathrin-mediated endocytosis (36), a key mechanism of viral entry. Given the limited studies on ADDA 5 in the context of viral infections, we decided to investigate its potential antiviral effects against HCoV-OC43. A dose–response curve in A549-ACE2 cells revealed a 50% cytotoxic concentration (CC_50_) of 33 μM for ADDA 5 (Fig. 2*H*, blue). Hence, we applied 20 μM (∼75% cell viability) to A549-ACE2 cells as a post-treatment following HCoV-OC43 infection. Notably, this treatment did not compromise the viability of infected cells (Fig. 2*H*, red and *SI Appendix*, Fig. S2*B*) while significantly reducing intracellular viral gene expression at 48 h post-infection (hpi) (Fig. 2*I*). The plaque assay further demonstrated a 2.2 log_10_ reduction (99.4%) in extracellular viral load following ADDA 5 treatment (Fig. 2*J* and *SI Appendix*, Fig. S2*C*), providing experimental evidence that CIV inhibition effectively blocks HCoV-OC43 replication. Taken together, our results confirm that the anti-HCoV targets identified by our framework can indeed suppress viral replication.

### Development of an scRNA-seq-Applicable Framework for HCoV-OC43-infected Human Organoids

Organoids more accurately recapitulate tissue architecture and intercellular interactions than conventional monolayer cultures. Recent advances in organoid technology and transcriptomic profiling have enabled detailed characterization of host responses across diverse cell types during viral infection (37, 38). In particular, respiratory organoids differentiated under air–liquid interface (ALI) conditions reproduce cellular exposure to air and serve as robust models for studying airborne pathogens (39, 40). However, the complexity and high dimensionality of scRNA-seq data make it challenging to systematically identify antiviral host factors. To address this limitation, we modified our computational framework to be compatible with scRNA-seq data and evaluated its ability to identify reliable antiviral targets. We used our previously published dataset GSE262439 (40), which profiled HBE organoids infected with four beta-coronaviruses (HCoV-OC43, SARS-CoV-2, MERS-CoV, and SARS-CoV) (Fig. 3*A*).

**Fig. 3.**
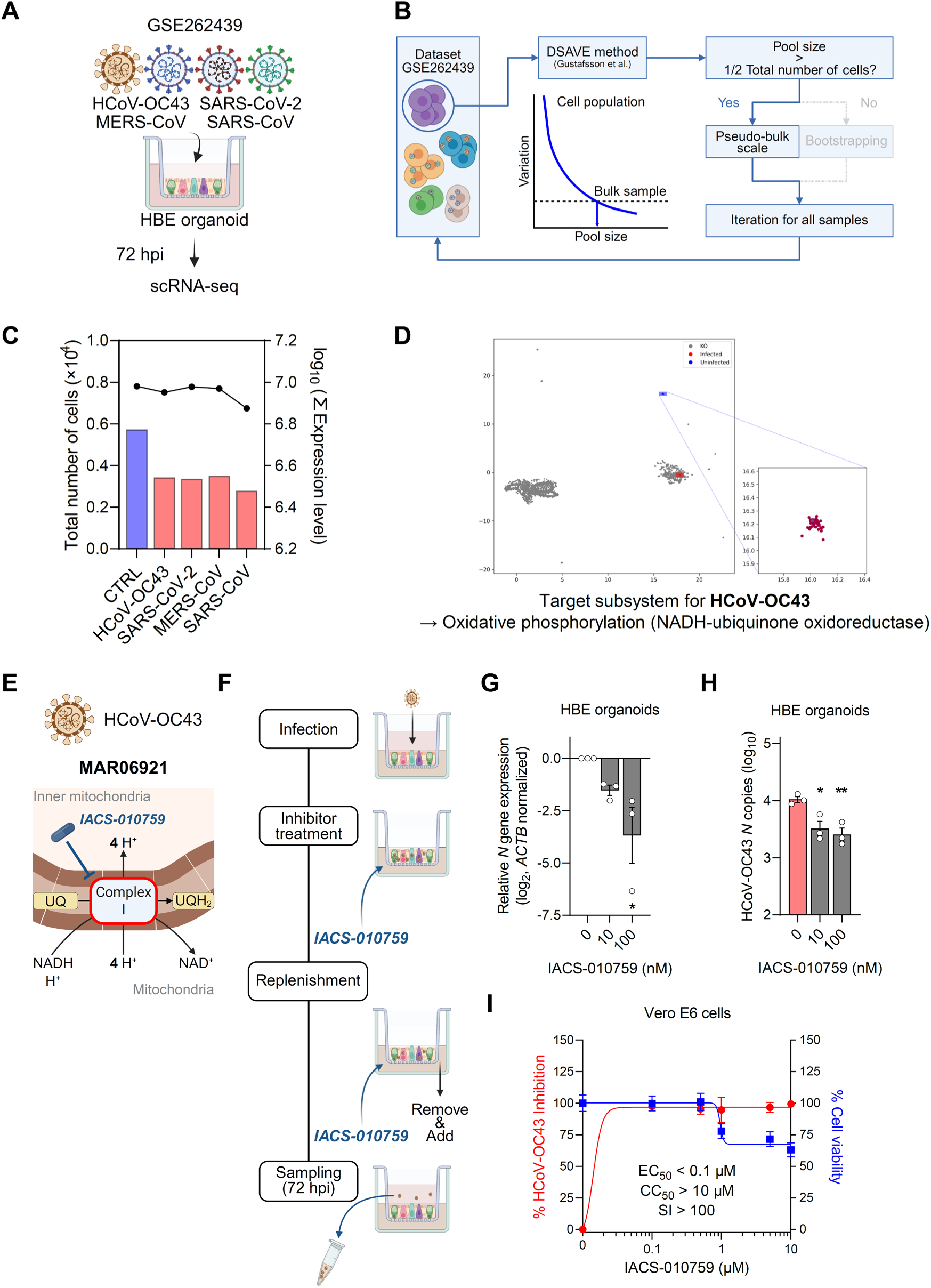
Application of the framework to HCoV-OC43–infected HBE organoids. **(A)** Graphical summary of the experimental design for the scRNA-seq dataset GSE262439. Images were created with BioRender.com. **(B)** Schematic flow of applying the DSAVE method. The cell pooling size (a threshold of CPM > 0.5) was estimated based on the total variation score. If the estimated pooling size did not exceed half of the total number of cells in a sample group, bootstrapping with random mixing was performed; otherwise, the entire sample was used at the pseudo-bulk scale. This procedure was repeated for each sample group individually. Images were created with BioRender.com. **(C)** Total number of cells (bars) and gene expression levels (points with connecting lines) for each sample group used in constructing infection-specific GEMs. Expression levels represent normalized RNA counts. **(D)** UMAP plots comparing fluxomes across all GEMs derived from the GSE262439 dataset for HCoV-OC43. The uninfected sample is represented in blue, the virus-infected sample in red, and the single-gene knockout simulation results as gray dots. A detailed gene list is provided in Table S2. **(E)** Schematic representation of the OXPHOS complex I (NADH–ubiquinone oxidoreductase) reaction and its inhibition by IACS-010759 (adapted from https://metabolicatlas.org/). Images were created with BioRender.com. UQ: ubiquinone, UQH_2_: ubiquinol. **(F)** Experimental design for evaluating the anti-HCoV-OC43 effect of IACS-010759 in HBE organoids (see *Materials and Methods* for details). Images were created with BioRender.com. **(G)** Intracellular HCoV-OC43 nucleocapsid (*N*) gene expression at 72 hpi was quantified via RT-qPCR (n = 3). *ACTB* was the normalization control. Statistical significance was calculated using one-way ANOVA with Dunnett’s multiple comparison test. **(H)** Extracellular HCoV-OC43 RNA copies at 72 hpi were quantified by RT-qPCR targeting nucleocapsid (*N*) amplicon. For each sample, 100 μl of viral supernatant was collected from infected HBE organoids (n = 3). Statistical significance was calculated using one-way ANOVA with Dunnett’s multiple comparison test. **(I)** Dose–response analysis for IACS-010759 in Vero E6 cells. Cell viability after 72 h of treatment (blue curve, n = 4) and HCoV-OC43 titer at 72 hpi (red curve, n = 4) were measured. Curves were fitted by nonlinear four-parameter sigmoidal regression in GraphPad Prism. HCoV-OC43 inhibition (%) was calculated relative to DMSO controls. SI: selectivity index.

To construct GEMs from scRNA-seq data, we first assessed the suitability of a bootstrapping strategy using the downsampling-based variation estimation (DSAVE) method (41) (Fig. 3*B*). DSAVE showed that achieving stable gene expression profiles with bias comparable to bulk RNA-seq data required pooling more than half of the cells per sample, rendering bootstrapping infeasible (*SI Appendix*, Fig. S3). Therefore, we reconstructed infection-specific GEMs at a pseudo-bulk scale using all cells from each sample. Although the total number of cells varied (2,781–5,727 cells), normalized read counts were comparable across samples, except for the SARS-CoV–infected organoids, which were excluded from further analysis (Fig. 3*C*).

As in the cell line models, we performed single-gene knockout simulations on the organoid GEMs and analyzed metabolic flux profiles from control, infected, and simulated knockout conditions using UMAP. This analysis again revealed antiviral targets capable of shifting infected metabolic states toward those of uninfected cells (Fig. 3*D*). Notably, the OXPHOS subsystem emerged again as a vulnerability in HCoV-OC43–infected organoids, consistent with our findings in monolayer cultures (Fig. 2*B*). In organoids, however, the predicted targets encompassed 44 subunits of OXPHOS complex I (*SI Appendix*, Table S3).

Complex I (NADH–ubiquinone oxidoreductase, CI) initiates the respiratory chain by oxidizing NADH and transferring electrons to ubiquinone (Fig. 3*E*). Among known CI inhibitors, rotenone is primarily used as an insecticide, while the FDA-approved type II diabetes drug metformin has been reported to exert broad-spectrum antiviral activity (42). We therefore selected IACS-010759, a selective CI inhibitor (43) without prior application in antiviral studies, for experimental validation. HBE organoids were infected with HCoV-OC43 via the apical side, and IACS-010759 was added daily to the basolateral compartment under ALI conditions (Fig. 3*F*). Consistent with computational predictions, treatment with IACS-010759 led to a dose–dependent reduction in viral gene expression (Fig. 3*G* and *SI Appendix*, Fig. S4*A*) and viral RNA copy number in the supernatant (Fig. 3*H* and *SI Appendix*, Fig. S4*B*). At 100 nM, both intracellular and extracellular viral RNA levels decreased, although mild cytotoxicity (< 75% viability) was observed (*SI Appendix*, Fig. S4*C*). To further confirm specificity, we tested IACS-010759 in Vero E6 cells, a standard model for coronavirus research. The drug exhibited potent antiviral activity at non-cytotoxic concentrations (below 0.5 μM) (Fig. 3*I*), yielding a high selectivity index (CC_50_/EC_50_) against HCoV-OC43. Together, these results validate OXPHOS complex I inhibition as an effective antiviral strategy identified by our framework, demonstrating that computationally derived targets can translate into experimentally verifiable therapeutic vulnerabilities.

### Extension of the Framework to Identify Antiviral Targets in SARS-CoV-2 and MERS-CoV

Encouraged by the successful validation of antiviral targets in HCoV-OC43–infected organoids, we applied our framework to two high-priority coronaviruses lacking effective treatments: SARS-CoV-2 and MERS-CoV. Computational analysis of scRNA-seq data from infected HBE organoids consistently identified genes associated with pyrimidine catabolism as candidate antiviral targets (Fig. 4*A* and *B* and *SI Appendix*, Table S4 and S5). Although annotated under both pyrimidine metabolism and β-alanine metabolism subsystems in Human1, the identified reactions converge on the pyrimidine catabolic pathway, hereafter referred to as “pyrimidine catabolism.” This pathway comprises a three-step enzymatic process converting uracil and thymine into β-alanine and D-3-aminoisobutanoate, respectively. The first and rate-limiting step is catalyzed by dihydropyrimidine dehydrogenase (DPYD), which reduces pyrimidines to dihydropyrimidines (Fig. 4*C*). Notably, DPYD has not been previously studied in the context of viral infection, prompting us to examine its potential as a therapeutic target.

**Fig. 4.**
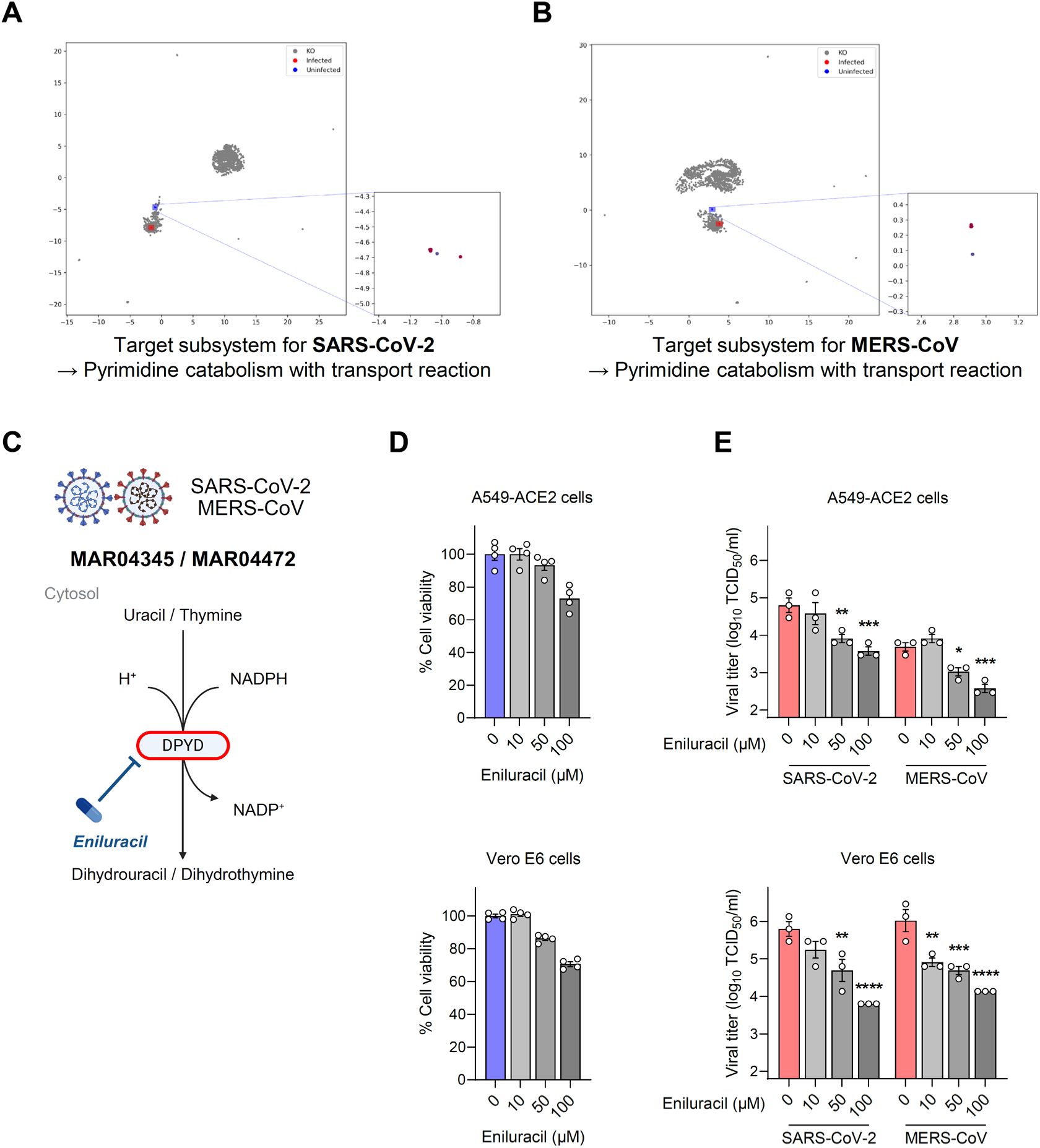
Application of the framework for SARS-CoV-2– and MERS-CoV–infected HBE organoids. **(A and B)** UMAP plots comparing fluxomes across all GEMs derived from the GSE262439 dataset for SARS-CoV-2 **(A)** and MERS-CoV **(B)**. The uninfected sample is represented in blue, the virus-infected sample in red, and the single-gene knockout simulation results as gray dots. Detailed gene lists are provided in Table S3. **(C)** Schematic representation of the reaction involving DPYD and its inhibition by Eniluracil (adapted from https://metabolicatlas.org/). Images were created with BioRender.com. **(D)** Dose–response analysis for Eniluracil in A549-ACE2 and Vero E6 cells. Cell viability was measured after 48 h of treatment (n = 4). **(E)** Extracellular SARS-CoV-2 and MERS-CoV titers at 48 hpi were measured by the endpoint dilution assay (n = 3). Statistical significance was calculated using two-way ANOVA with Dunnett’s multiple comparison test.

We first evaluated the cytotoxicity of Eniluracil, a small chemical inhibitor of DPYD, in A549-ACE2 and Vero E6 cells (Fig. 4*D*). Both cell types tolerated concentrations up to 50 μM without notable cytotoxicity, while mild toxicity was observed at 100 μM. In A549-ACE2 cells, a significant reduction in viral RNA copy number was detected only at 100 μM (*SI Appendix*, Fig. S5*A*). Nevertheless, viral titers decreased substantially at 50 μM, with reductions of 87.0% for SARS-CoV-2 and 78.5% for MERS-CoV (Fig. 4*E*). In Vero E6 cells, the Eniluracil treatment at 50 μM robustly inhibited replication of both viruses (92.3% for SARS-CoV-2 and 95.4% for MERS-CoV), and notably, MERS-CoV replication was suppressed by over 90% even at 10 μM, corresponding to a favorable selectivity index (> 10) (Fig. 4*E*). Reductions in viral RNA copy numbers further corroborated these results (*SI Appendix*, Fig. S5*B*). These findings establish DPYD inhibition and possibly pyrimidine catabolism as a previously unrecognized vulnerability in SARS-CoV-2 and MERS-CoV, extending the applicability of our framework to uncover novel, host-directed antiviral (HDA) strategies.

To test whether this antiviral effect extends to more physiologically relevant systems, we evaluated Eniluracil in SARS-CoV-2– and MERS-CoV–infected HBE organoids. Eniluracil exhibited no detectable cytotoxicity at concentrations up to 100 μM (*SI Appendix*, Fig. S6*A*). Because drug penetration can be limited in three-dimensional structures, we modified the treatment protocol by administering Eniluracil to both the basolateral and apical compartments(44) (Fig. 5*A*). Under these optimized conditions, immunohistochemistry (IHC) revealed a pronounced reduction of viral nucleocapsid protein levels in Eniluracil-treated organoids, indicating suppressed replication (Fig. 5*B* and *C*). Consistently, viral titers and RNA copy numbers in the supernatant were significantly decreased (Fig. 5*D* and *SI Appendix*, Fig. S6*B*), confirming the antiviral efficacy of Eniluracil against both SARS-CoV-2 and MERS-CoV in the organoid model. Importantly, gene expression profiles and pathway enrichment analyses from scRNA-seq data showed no significant alterations in OXPHOS or pyrimidine catabolism pathways (*SI Appendix*, Fig. S7*A*–*E*), underscoring that our computational framework can uncover therapeutic vulnerabilities not captured by conventional transcriptomic analyses.

**Fig. 5.**
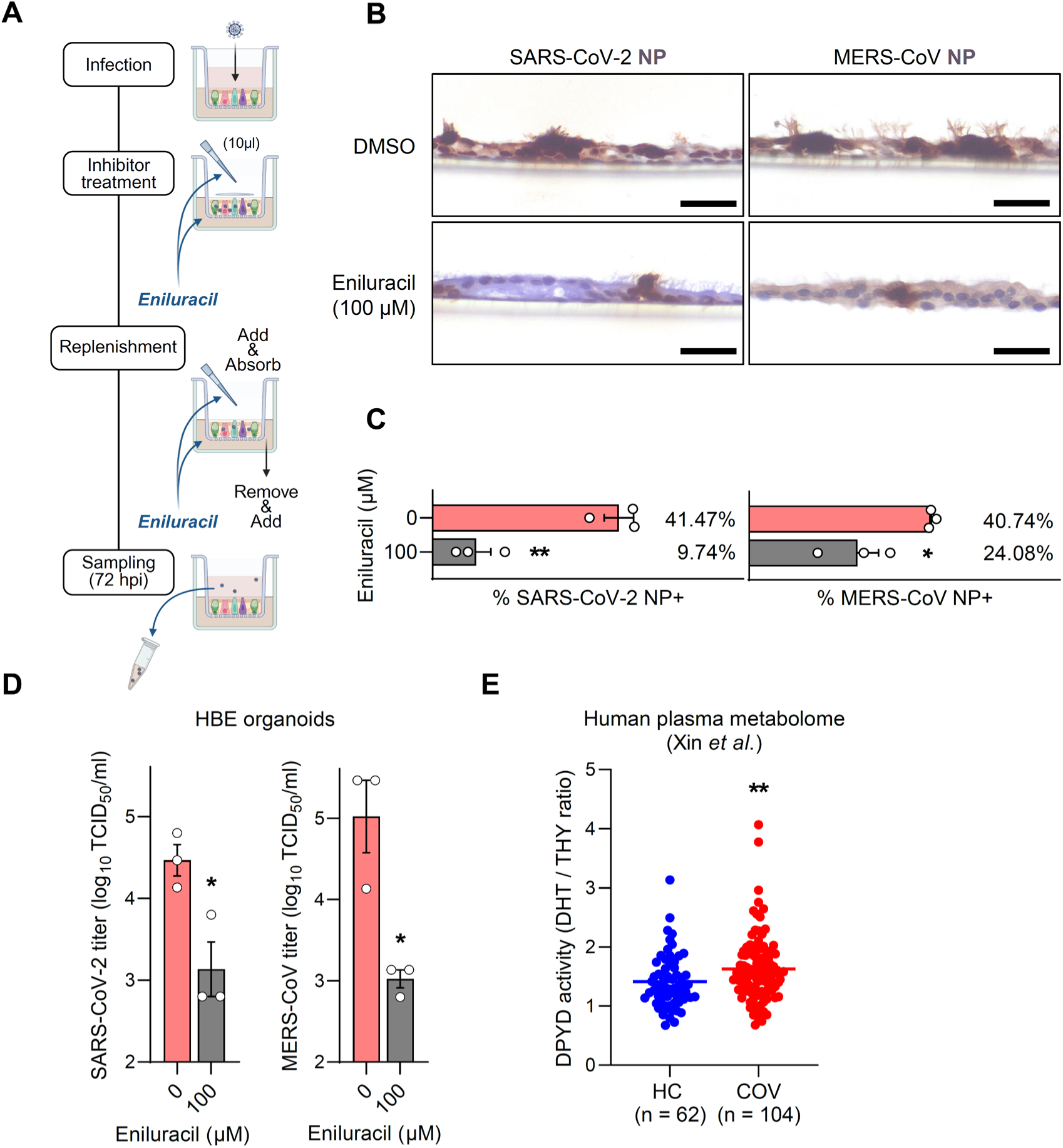
Effects of DPYD inhibition on SARS-CoV-2 and MERS-CoV replication in organoids. **(A)** Experimental design for evaluating the antiviral effects of Eniluracil against SARS-CoV-2 and MERS-CoV in HBE organoids (see *Materials and Methods* for details). Images were created with BioRender.com. **(B)** Representative IHC images of organoid sections. Viral nucleocapsid (NP) was used as the infection marker for each virus. Scale bar = 20 μm. **(C)** Quantification of NP-positive (NP+) organoids (n = 3). The NP+ fraction was calculated as NP+ area divided by total cellular area, with areas measured using QuPath software. Statistical significance was calculated using a one-tailed Student’s t-test. **(D)** Extracellular SARS-CoV-2 and MERS-CoV titers at 72 hpi were measured by the endpoint dilution assay (n = 3). Statistical significance was calculated using a one-tailed Student’s t-test. **(E)** Re-analysis of a human plasma metabolome dataset (n = 166). DPYD enzymatic activity was calculated as the ratio of DHT to THY (raw intensities) for each sample (HC: healthy control, COV: COVID-19 patient). Statistical significance was calculated using a two-tailed Welch’s t-test.

To further assess the clinical potential of targeting DPYD, we analyzed its enzymatic activity in patients with COVID-19. DPYD activity was estimated using the metabolite ratio of dihydrothymine (DHT) to thymine (THY) (45, 46). Re-analysis of a publicly available plasma metabolome dataset (47) revealed a significant increase in the [DHT] to [THY] ratio in patients with COVID-19 (n = 104) compared with healthy controls (n = 62) (Fig. 5*E*). These findings indicate the enhanced DPYD activity during SARS-CoV-2 infection, supporting its potential as a clinically relevant antiviral target.

## Discussion

In this study, we developed and experimentally validated a computational framework that combines GEMs with single-gene knockout simulation followed by UMAP-based dimensionality reduction for rapid discovery of antiviral targets. The main idea of our framework is “metabolic rewiring,” which involves identifying metabolic enzymes whose inhibition can restore virus-altered metabolic profiles to a state resembling the pre-infection condition (48–50). To this end, we meticulously constructed infection-specific GEMs based on transcriptomic data and analyzed single-gene knockout simulations based on these GEMs via UMAP to effectively pinpoint key target genes among the various metabolic pathways changed during viral infection. The feasibility of this approach was previously demonstrated in a drug-resistant breast cancer cell model (30).

Applying our framework to MRC-5 cells infected with HCoV-229E identified genes associated with porphyrin metabolism. Although elevated serum porphyrin levels have been reported in patients with hepatitis C virus, human immunodeficiency virus, and SARS-CoV-2 (51, 52), the relationship between host porphyrin and viral infection remains unclear. Moreover, the antiviral activity of suppressing porphyrin metabolism was unexplored. In the case of A549-ACE2 cells infected with HCoV-OC43, the OXPHOS subsystem (complex IV) emerged as an antiviral target. Viral modulation of host OXPHOS pathways is known to vary depending on both virus species and host cell types (53, 54). While several OXPHOS inhibitors have shown antiviral activity against HCoVs (27, 55–58), their precise mechanisms of action are difficult to delineate due to the close metabolic coupling between OXPHOS and the TCA cycle (43, 59). Moreover, recent virus-host interactome studies suggest potential interactions between host OXPHOS components and viral nonstructural proteins (60–62). Hence, more comprehensive mechanistic studies from both viral and host perspectives are needed to clarify how the predicted targets can suppress HCoV replication.

In HBE organoids, the OXPHOS subsystem again emerged as a key antiviral target for HCoV-OC43, but corresponding to complex I rather than complex IV. This discrepancy possibly reflects the concept of our framework, which prioritizes gene-level regulators capable of optimally rewiring the host metabolic network through GPR association, rather than relying solely on the magnitude of differential fluxes. Indeed, OXPHOS complex I (MAR06921) also showed a marked flux increase in HCoV-OC43–infected A549-ACE2 cells (Fig. 2*C*), yet were not nominated as targets. Extension of the framework to SARS-CoV-2– and MERS-CoV–infected organoids revealed the pyrimidine catabolism as a common vulnerability. While the antiviral effects of enzymes involved in *de novo* pyrimidine synthesis are well documented (7, 63, 64), to the best of our knowledge, this is the first study to implicate the pyrimidine catabolic process in viral infection. Moreover, such key metabolic pathways could not be discovered under conventional transcriptomic analyses, including differential gene expression or pathway enrichment analyses (*SI Appendix*, Fig. S7*C*–*E*). These findings collectively highlight the advantage of our framework in providing unique, unbiased insights without prior knowledge of the virus.

It is notable that our framework highlights genes encoding solute carrier proteins (SLCs) involved in transport reactions as potential antiviral targets. The SLC family comprises ∼450 membrane transporters that mediate the exchange of diverse substrates (65, 66), a subset of which can act as upstream cues of metabolic pathways (67–69). We showed that knockdown of the glycine transporter gene *SLC25A38*, participating in the early steps of porphyrin metabolism, effectively reduced HCoV-229E replication. Analysis of the HBE organoid scRNA-seq dataset further revealed the *SLC32A1* gene, annotated as an extracellular transport of glycine and β-alanine, as a common candidate target for SARS-CoV-2 and MERS-CoV (*SI Appendix*, Table S4 and S5). In addition to our findings, several studies also reported the involvement of the SLCs in viral infections, such as bile salt transporter SLC10A1 (NTCP) for hepatitis B virus (70), amino acid transporter SLC3A2 (CD98 heavy chain) for hepatitis C virus (71), sialic acid transporter SLC35A1 for vesicular stomatitis virus (72), and arginine sensor SLC38A9 for SARS-CoV-2 (73). Taken together, we propose that certain SLCs may serve as key regulators of host metabolic rewiring in response to viral infections. Based on previous studies, it would be valuable to investigate how perturbation of SLC activity influences global host metabolism. Further systematic studies on SLC transporters could clarify their roles in viral replication and explore their therapeutic potential as effective antiviral targets.

One limitation of our study is that some of the identified targets may encode key regulators of host metabolism that are closely linked to cell survival. For example, the OXPHOS subsystem, identified from HCoV-OC43–infected samples, is a central pathway for host energy and redox homeostasis. While targeting this pathway can suppress viral replication, it may also compromise the viability of host cells. Therefore, antiviral strategies that interfere with such essential pathways should be approached with caution, and the trade-off between antiviral efficacy and host cell toxicity must be carefully considered. Nonetheless, we were able to determine the concentration of the OXPHOS inhibitor that yields antiviral effects without significant cytotoxicity. Another limitation of such HDAs is the difficulty in excluding potential off-target effects. Although we utilized selective inhibitors with well-established metabolic functions, they may induce unexpected effects on cell signaling pathways by inhibiting off-target enzymes. Nevertheless, HDAs offer several advantages, including a lower risk of resistance, applicability across diverse variants, and feasibility for drug repurposing, making them a promising strategy against future viral threats (74–76).

In summary, our computational framework enables the detection of previously overlooked metabolic targets by focusing on virus-induced alterations in host metabolism. The experimental validation of these targets underscores the robustness of our approach, suggesting its potential applicability across a broad spectrum of virus-infected samples. We envision the developed framework serving as a strategic foundation for exploring tailored HDAs against emerging and recurrent viruses.

## Materials and Methods

### Infection-specific GEM extraction, flux simulation, and dimensionality reduction

Infection-specific GEMs were constructed for control and infected samples from each dataset using the Human1 template (77) (version 1.6.0) and rank-based task-driven integrative network inference for tissues (rank-based tINIT) (78). Metabolic fluxes for both control and infected samples were calculated using the least absolute deviation (LAD) method (29), with a threshold of 0.01 imposed as a constraint on the biomass formation rate to simulate a more realistic flux state (i.e., a cell growth rate > 0 for the control sample). For single-gene knockout simulation of the infected model, flux changes were predicted using linear minimization of metabolic adjustment (lMOMA) (79). In this case, constraints were introduced by setting the upper and lower bounds of reactions eliminated by single-gene knockout to 0. Reconstruction of all infection-specific GEMs was performed in MATLAB (R2021a) using Gurobi Optimizer (version 9.1.2). Reading, writing, and manipulation of COBRA-compliant MATLAB files were implemented using COBRApy (version 0.23.1) (18). Dimensionality reduction analysis of total flux sets (control samples, infected samples, and single-gene knockout simulation results) was performed using UMAP (version 0.5.1) (80) with optimized parameters (n_neighbors = 3 or 7, min_dist = 0.1). All UMAP plots were visualized using Seaborn (version 0.11.0) and Matplotlib (version 3.3.4) libraries. Target genes whose knockout flux profiles clustered with the control sample were identified using density-based spatial clustering of applications with noise (DBSCAN) with parameters eps = 0.3 and min_samples = 3 (81).

### RNA-seq data processing

A total of three publicly available RNA-seq datasets were utilized in this study (details summarized in *SI Appendix*, Table S6). To construct infection-specific GEMs, we used averaged transcripts per million (TPM) values across 3 biological replicates. When datasets provided only raw read counts, transcript lengths were retrieved from the Ensembl API (82), followed by TPM normalization. Based on raw read counts, DEGs were identified using the DEGseq package (version 1.14.0) (83) for bulk RNA-seq and the DESeq2 (version 1.46.0) (84) for scRNA-seq. GO enrichment analysis and GSEA were performed with the clusterProfiler package (version 4.14.3) (85). Results were visualized using the ggplot2 (version 3.5.1) and ComplexHeatmap (version 2.22.0) packages (86) in R (version 4.4.2).

### Viral titration

Human beta-coronavirus strain OC43 (HCoV-OC43) and alpha-coronavirus strain 229E (HCoV-229E) were purchased from the Korea Bank for Pathogenic Viruses (KBPV, Republic of Korea). For HCoV-OC43, its propagation and titration were performed as previously described (87) with an additional endpoint dilution assay using MRC-5 lung fibroblast cells at 5 days post-infection. HCoV-229E stocks were stored at −80 °C without further propagation, and titers were determined in MRC-5 cells. SARS-CoV-2 (Omicron BA.2 variant) and MERS-CoV (strain KOR/KNIH/2015) stocks were obtained from the Virus Research Resource Center (VRRC) at the Institute for Basic Science. Viral titers were determined in Vero E6 cells by assessing cytopathic effects at 3 days post-infection and calculating TCID_50_/ml by the Spearman–Kärber method.

### Small molecule inhibitor and cytotoxicity assay

IACS-010759, ADDA 5 hydrochloride, and Eniluracil were purchased from MedChemExpress, USA, and were dissolved in DMSO following the manufacturer’s instructions. The cytotoxicity of each inhibitor was evaluated in A549-ACE2 or Vero E6 cells using the CellTiter-Glo Luminescent Cell Viability Assay (Promega, USA). For HBE organoids, inhibitors were added to the basolateral side and replenished daily. Organoid viability was measured 72 h after the treatment using the CellTiter-Glo 3D Cell Viability Assay (Promega, USA) following the provided manual. Luminescence signals were recorded with a multimode microplate reader. Changes in cell viability upon ADDA 5 treatment in HCoV-OC43–infected cells were assessed using the Sulforhodamine B assay, as previously described (88).

### Generation of shRNA-mediated knockdown cells

shRNA oligos targeting *ALAD* and *SLC25A38* were designed from the GPP web portal (Broad Institute) with the highest adjusted score. The hairpin sequences for each target are listed in Table S7. For shRNA subcloning, the pLKO.1-TRC cloning vector (Addgene, USA) plasmid was used as the template. The vector was digested using the restriction enzymes AgeI-HF and EcoRI-HF (New England Biolabs, USA). The digested vector was separated by gel electrophoresis, followed by gel extraction using the Expin Gel SV kit (GeneAll, Republic of Korea). The shRNA oligomers were annealed at 95 °C for 1 min and subsequently ligated together with the digested vector and T4 DNA ligase (New England Biolabs, USA). Ligated plasmids were then transformed into DH5α chemically competent *E. coli* (Enzynomics, Republic of Korea) through heat shock transformation. After 2 days of ampicillin selection, the shRNA plasmid vectors were purified using the NucleoBond Xtra-Midi kit (Macherey-Nagel, Germany). All shRNA subcloning results were validated by difficult Sanger sequencing (Macrogen, Republic of Korea). For the control group, the pLKO.1-scramble shRNA (Addgene, USA) plasmid was processed in the same manner to construct the shScramble plasmid vector.

shRNA-containing lentiviral vectors (LVVs) for control (Scramble) and targeting *ALAD* and *SLC25A38* mRNAs were generated following a previously published protocol (89). LVVs were then stored at −80 °C and not subjected to multiple freeze-thaw cycles to prevent titer reduction. To knock down the target gene expression, 2.0×10^5^ cells were reverse transduced with 100 μl LVVs and 10 ng/ml polybrene in 12-well plates. This was followed by 5 days of puromycin selection to obtain the knockdown cells.

### Validation of antiviral targets in cell lines

For HCoV-229E, 2.0×10^5^ knockdown cells were plated in a 12-well plate without puromycin. After a 1-day recovery period, cells were infected at a multiplicity of infection (MOI) of 0.01 for 1.5 h at 35°C, followed by a DPBS wash to remove the remaining virus. At 24 hpi, supernatant and cell lysate were collected. The parallel experiment was repeated with MRC-5 WT cells.

For HCoV-OC43, 4.0×10^5^ A549-ACE2 cells were seeded in a 6-well plate and maintained at 37 °C with 5% CO_2_ for 24 h. Cells were infected with the virus at an MOI of 1 for 1 h at 35°C. Following infection, cells were rinsed again and replenished with DMEM (Welgene, Republic of Korea) supplemented with 5% FBS (Gibco, USA), and either DMSO or 20 μM ADDA 5 inhibitor. To assess the impact of the ADDA 5 inhibitor on viral gene expression, total RNA was extracted from cell lysates, and supernatants were collected at 48 hpi.

For SARS-CoV-2 and MERS-CoV, dose–response experiments were carried out using 8.0×10^4^ A549-ACE2 or 5.0×10^4^ Vero E6 cells seeded in 24-well plates. In the BSL-3 facility, cells were infected at an MOI of 0.01 for 1 h at 37°C, washed with DPBS, and then incubated either in growth medium with DMSO or Eniluracil at concentrations ranging from 10 to 100 μM. At 48 hpi, the supernatant was harvested for viral RNA quantification and titration.

### Validation of antiviral targets in HBE organoids

HBE cells (ScienCell, USA) were expanded in PneumaCult-Ex Plus medium (STEMCELL Technologies, Canada) in a 150 mm cell culture dish. For organoid differentiation under ALI conditions, 8.5×10^4^ cells were seeded onto 12-well Transwell plates (Corning, USA). After 2 or 3 days, the apical medium was removed (air-lifted) and the basal medium was replaced with PneumaCult-ALI medium (STEMCELL Technologies, Canada). Cells were maintained under ALI conditions for 4 weeks to induce differentiation.

Differentiated HBE organoids were pre-rinsed with DPBS and exposed to viral inoculum for 2 h at 37°C. After infection, the inoculum was removed, and the apical side was washed twice with DPBS. For HCoV-OC43, either DMSO or IACS-010759 (10 or 100 nM) was supplied daily to the basal chamber medium. For SARS-CoV-2 and MERS-CoV, DMSO or 100 μM Eniluracil was replenished daily to the basal chamber, together with an additional 10 µL solution applied directly to the apical surface by pipetting. The supernatant was collected at 72 hpi for viral RNA quantification and titration. The remaining organoids were treated with TRIzol (for HCoV-OC43) or fixed with 4% paraformaldehyde (for SARS-CoV-2 and MERS-CoV) for further analyses.

### RNA extraction and reverse transcription-quantitative PCR

Total RNA was extracted using TRIzol (Invitrogen, USA) and precipitated with sodium acetate (Invitrogen, USA) and 2-propanol (Sigma-Aldrich, USA). The isolated RNA was treated with DNase I (Takara, Japan), and phase separation was induced using acid-phenol:chloroform, pH 4.5 (Thermo Fisher Scientific, USA). For cDNA synthesis, purified RNA was reverse transcribed using RevertAid reverse transcriptase (Thermo Fisher Scientific, USA). The resulting cDNA was then amplified using RealFast SYBR Kit (Geneer, Republic of Korea), and RT-qPCR was conducted with the QuantStudio 1 Real-Time PCR system (Applied Biosystems, USA). Primers used in this study are listed in Table S8.

### Immunohistochemistry

Immunohistochemical staining of paraffin-embedded HBE organoids was conducted on a BOND RX instrument (Leica, Germany) according to the manufacturer’s protocols. Sections were incubated for 60 min at room temperature with either an anti-SARS-CoV-2 nucleocapsid antibody (Abcam, Cat # ab271180) or an anti-MERS-CoV nucleocapsid antibody (SinoBiological, Cat # 400680-MM10). Detection and visualization of viral nucleocapsid protein (NP) were performed using the Bond Polymer Refine Detection kit (Leica, Cat # DS9800), which includes the secondary antibody, hematoxylin counterstain, and DAB (3,3’-diaminobenzidine) chromogen. Viral NP-positive organoids were quantified using QuPath (v0.5.1).

### Statistical analysis

Statistical analyses were conducted using GraphPad Prism (version 10.2.2). Statistical details of experiments and analyses are described in the figure legends. Error bars represent the standard error of the mean. A p-value < 0.05 was considered statistically significant (* p < 0.05; ** p < 0.01; *** p < 0.001; **** p < 0.0001; n.s., not significant).

## Acknowledgments

We thank all members of the Center for Study of Emerging and Re-emerging Viruses in Korea Virus Research Institute for their helpful discussions and feedback on this paper. We also thank all members of the Hyun Uk Kim and Yoosik Kim laboratories.

## Data and materials availability

All data needed to evaluate the conclusions in the paper are present in the paper or the Supplementary Materials. Any additional information required to reanalyze the data reported in this paper is available from the corresponding author upon request.

