## Supplementary material for "Rapid discovery of antiviral targets through dimensionality reduction of genome-scale metabolic models": SI Appendix

**This PDF file includes:**

Figures S1 to S7
Tables S1 to S8

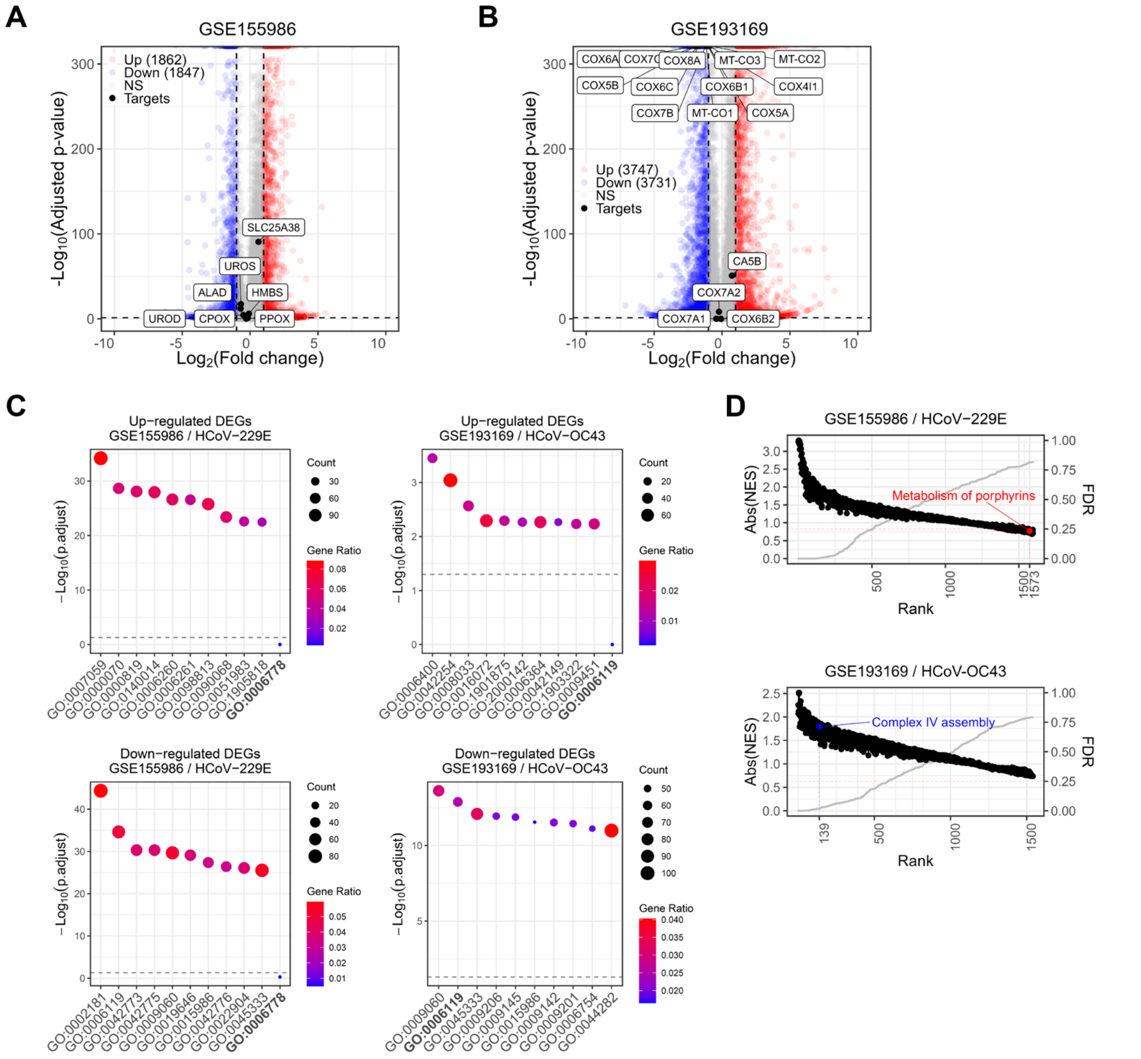


**Fig. S1. Comparison with conventional transcriptome analysis.**

**(A and B)** Volcano plots of DEGs in HCoV-229E–infected **(A)** and HCoV-OC43–infected **(B)** samples. Significantly upregulated genes (p-value < 0.05 and log2(fold change) > 1) are shown in red, and significantly downregulated genes (p-value < 0.05 and log2(fold change) < -1) are shown in blue. Antiviral target genes identified from our computational framework are shown in black with annotation. **(C)** Dot plots of GO enrichment results for significantly upregulated (p-value < 0.05 and log2(fold change) > 1) and downregulated (p-value < 0.05 and log2(fold change) < -1) DEGs. The top 10 biological processes are listed in descending order of adjusted p-value. Target subsystems (GO:0006778, metabolism of porphyrin-containing compounds; GO:0006119, oxidative phosphorylation) are highlighted in bold. **(D)** Rank plots of absolute normalized enrichment scores (NESs) for Reactome pathways identified by GSEA (p-value cutoff = 0.99). Gray dots indicate the false discovery rate (FDR) for each pathway. Target subsystems (Metabolism of porphyrins, R-HAS-189445; Complex IV assembly, R-HAS-9864848) are indicated.

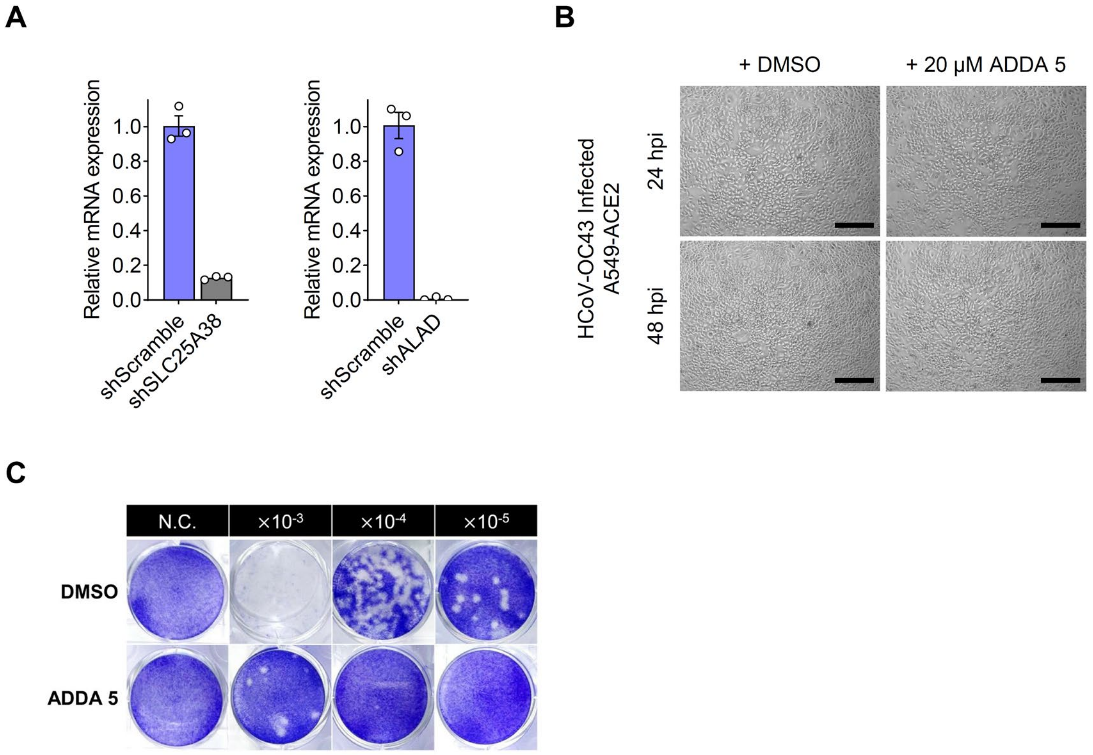


**Fig. S2. Validation of antiviral targets for HCoV-229E and HCoV-OC43.**
**(A)** The efficiency of shRNA-mediated gene knockdown was assessed using RT-qPCR (n = 3). Levels of *SLC25A38* or *ALAD* mRNAs in MRC-5 cells transduced with shScramble served as a control. The average of three biological replicates is shown with error bars denoting standard error of the mean (s.e.m). **(B)** Representative phase-contrast microscopy images of HCoV-OC43–infected A549-ACE2 cells. Scale bar = 200 µm. **(C)** A representative image of the HCoV-OC43 plaque assay performed in RD cells under control (DMSO) or ADDA 5-treated conditions.

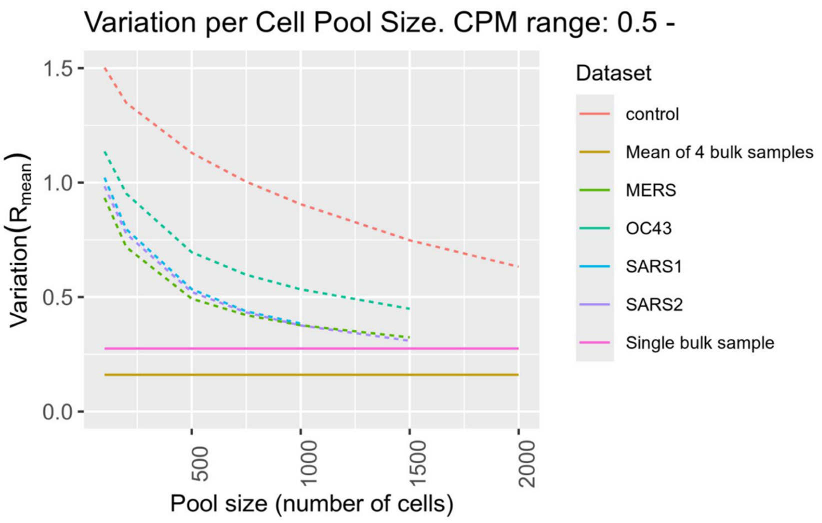


**Fig. S3. DSAVE-based estimation of total cell pool variation in the GSE262439 dataset.** Total variation scores were calculated using the DSAVE method to estimate the required pooling size of cells in the GSE262439 scRNA-seq dataset. The counts per million (CPM) threshold of 0.5 was applied.

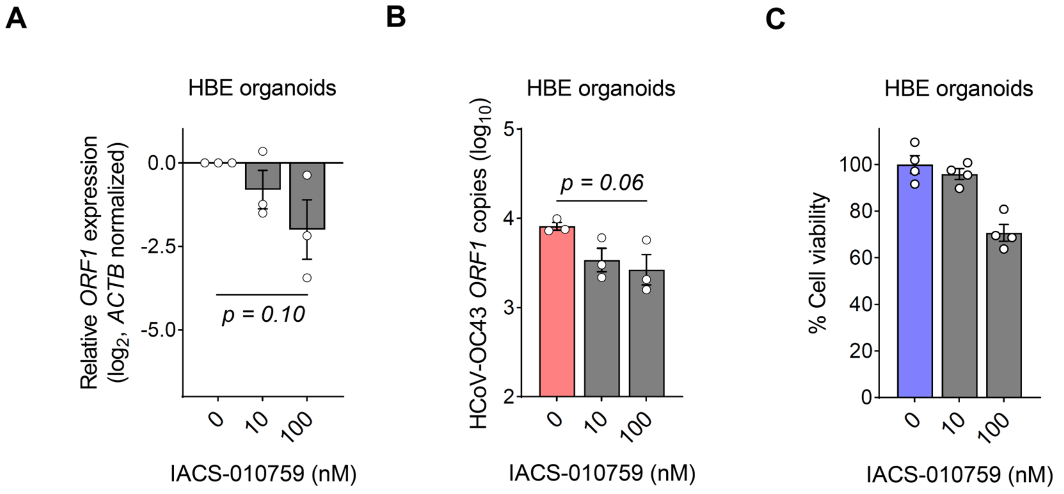


**Fig. S4. Effect of IACS-010759 on HCoV-OC43–infected HBE organoids.**
**(A)** Intracellular HCoV-OC43 open reading frame 1 (*ORF1*) gene expression at 72 hpi was quantified via RT-qPCR (n = 3). *ACTB* was used as the normalization control. **(B)** Extracellular HCoV-OC43 RNA copies at 72 hpi were quantified by RT-qPCR targeting the *ORF1* amplicon. For each sample, 100 µl of viral supernatant was collected from infected HBE organoids (n = 3). For **(A)** and **(B)**, the average of three biological replicates is shown with error bars denoting s.e.m. Statistical significance was calculated using one-way ANOVA with Dunnett's multiple comparison test. **(C)** Cell viability of HBE organoids treated with different concentrations of IACS-010759 was measured by CellTiter-Glo 3D Cell Viability Assay (n = 4). The average of biological replicates is shown with error bars denoting s.e.m.

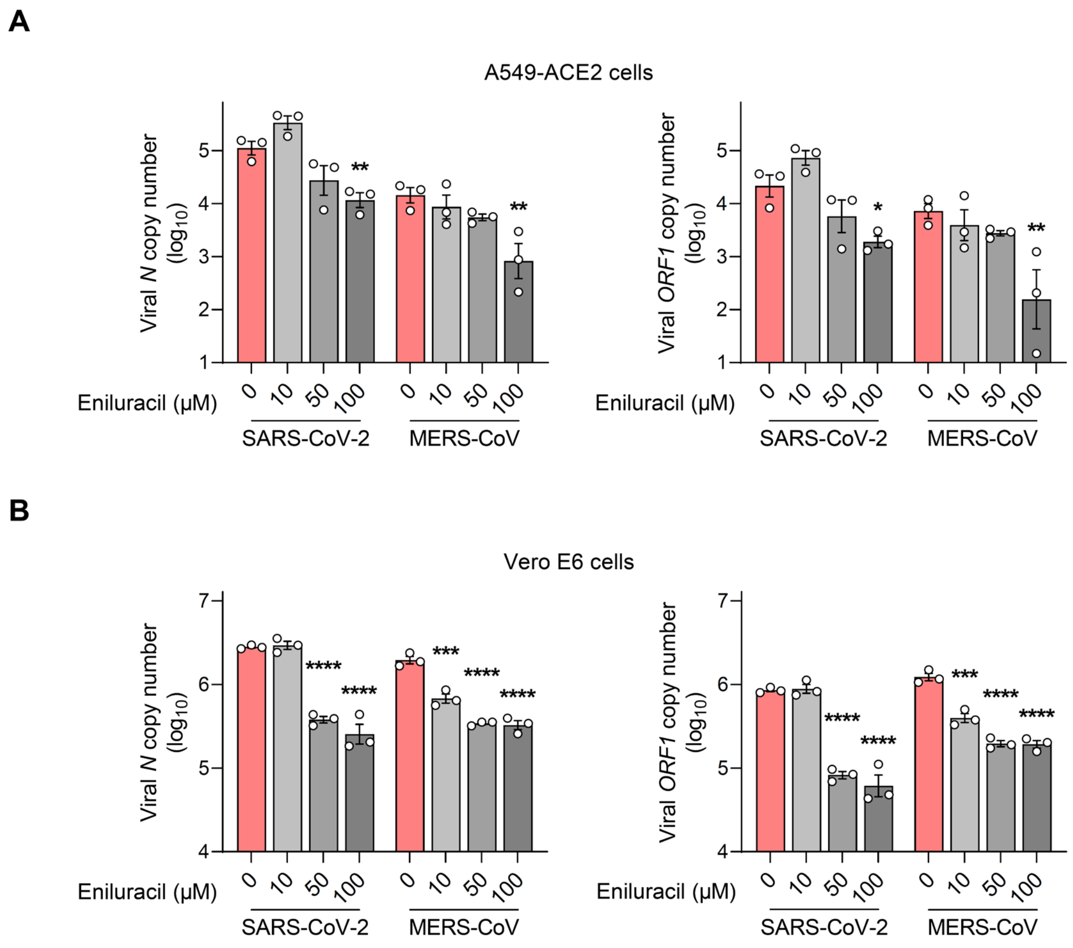


**Fig. S5. Effect of Eniluracil on SARS-CoV-2 and MERS-CoV replication in monolayer culture.**
**(A and B)** Extracellular SARS-CoV-2 and MERS-CoV RNA copies at 48 hpi were quantified by RT-qPCR targeting *N* and *ORF1* amplicons. For each sample, 100 µl of viral supernatant was collected from infected A549-ACE2 **(A)** and Vero E6 **(B)** cells. The average of three biological replicates is shown with error bars denoting s.e.m. Statistical significance was calculated using two-way ANOVA with Dunnett's multiple comparison test.

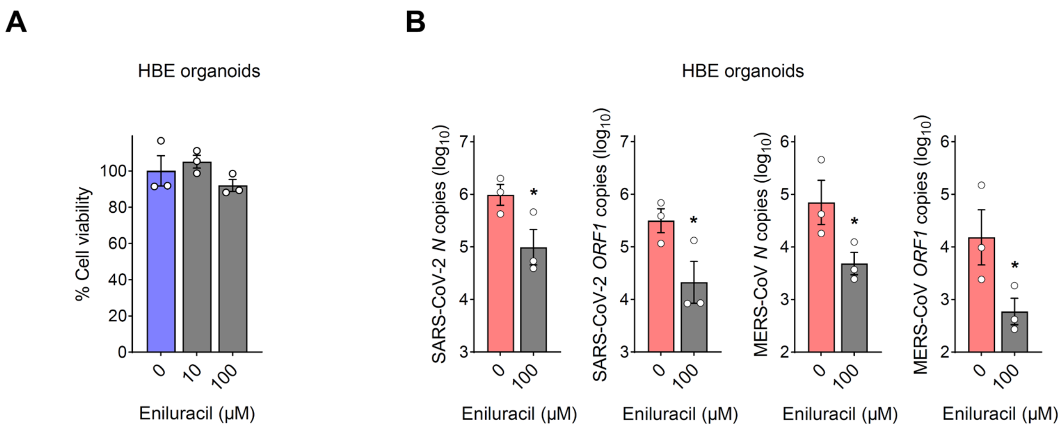


**Fig. S6. Effect of Eniluracil on SARS-CoV-2 and MERS-CoV replication in HBE organoids.** **(A)** Dose–response analysis for Eniluracil in HBE organoids. Cell viability was measured after 72 h of treatment. **(B)** Extracellular SARS-CoV-2 and MERS-CoV RNA copies at 72 hpi were quantified by RT-qPCR targeting *N* and *ORF1* amplicons. For each sample, 100 µl of viral supernatant was collected from infected HBE organoids. The average of three biological replicates is shown with error bars denoting s.e.m. Statistical significance was calculated using a one-tailed Student's t-test.

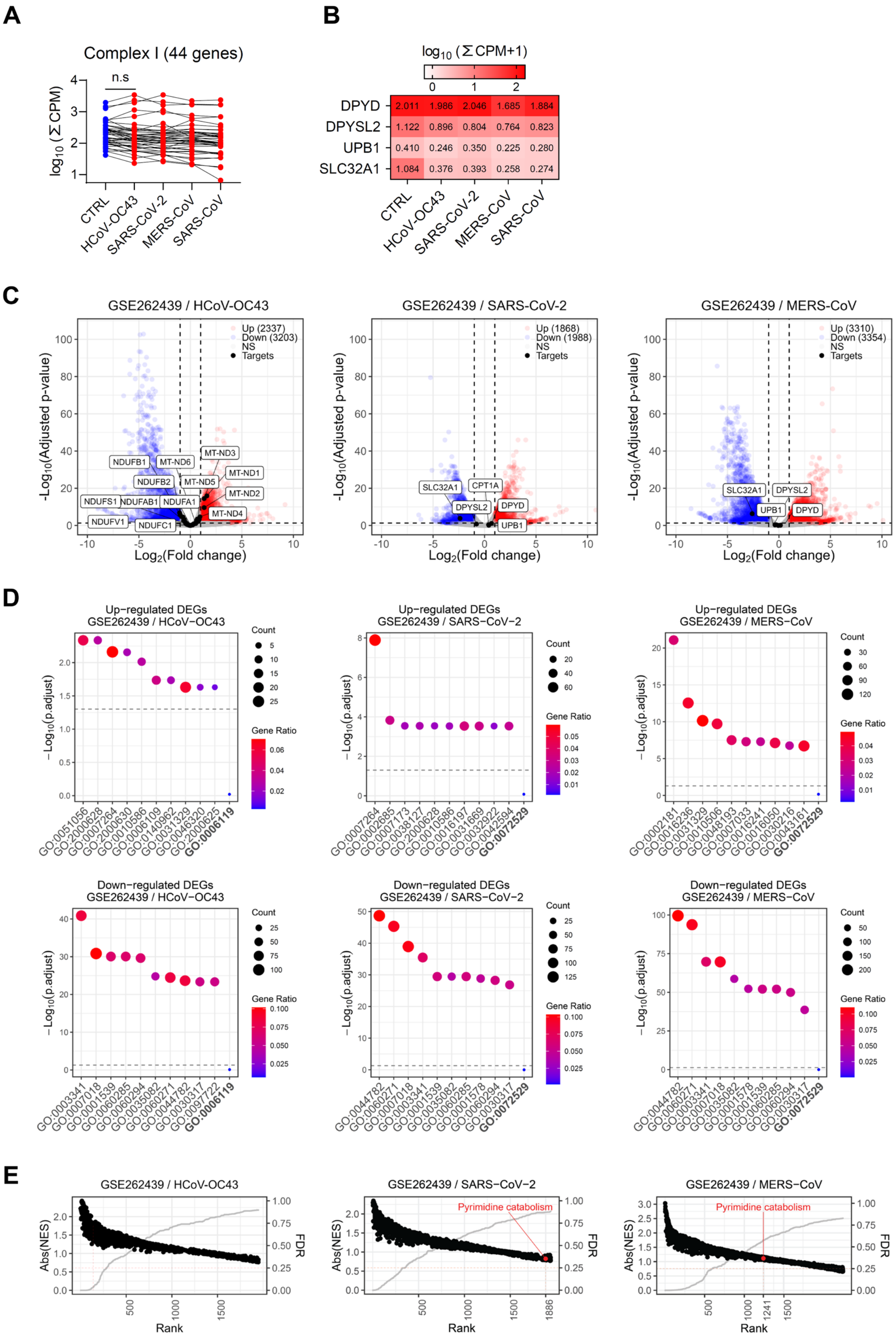
A
Complex I (44 genes)
n.s
log10 (Σ CPM)
CTRL
HCoV-OC43
SARS-CoV-2
MERS-CoV
SARS-CoV
B
log10 (Σ CPM+1)
DPYD 2.011 1.986 2.046 1.685 1.884
DPYSL2 1.122 0.896 0.804 0.764 0.823
UPB1 0.410 0.246 0.350 0.225 0.280
SLC32A1 1.084 0.376 0.393 0.258 0.274
CTRL
HCoV-OC43
SARS-CoV-2
MERS-CoV
SARS-CoV
C
GSE262439 / HCoV-OC43
Up (2337)
Down (3203)
NS
Targets
NDUFB1
MT-ND6
MT-ND3
NDUFB2
MT-ND5
MT-ND1
NDUFS1
NDUFAB1
NDUFA1
MT-ND2
NDUFV1
NDUFC1
MT-ND4
GSE262439 / SARS-CoV-2
Up (1868)
Down (1988)
NS
Targets
SLC32A1
CPT1A
DPYSL2
DPYD
UPB1
GSE262439 / MERS-CoV
Up (3310)
Down (3354)
NS
Targets
SLC32A1
DPYSL2
UPB1
DPYD
-Log10(Adjusted p-value)
Log2(Fold change)
D
Up−regulated DEGs
GSE262439 / HCoV−OC43
GSE262439 / SARS−CoV−2
GSE262439 / MERS−CoV
Down−regulated DEGs
GSE262439 / HCoV−OC43
GSE262439 / SARS−CoV−2
GSE262439 / MERS−CoV
Count
Gene Ratio
−Log10(p.adjust)
GO:0006119
GO:0072529
E
GSE262439 / HCoV-OC43
GSE262439 / SARS-CoV-2
GSE262439 / MERS-CoV
Pyrimidine catabolism
Abs(NES)
FDR
Rank
1886
1241

**Fig. S7. Comparison with conventional single-cell transcriptome analysis.**
**(A and B)** Distribution of total CPM per sample for 44 genes associated with OXPHOS complex I **(A)** and for 4 genes involved in pyrimidine catabolism **(B)**. Statistical significance was assessed using one-way ANOVA with Dunnett's multiple comparison test. **(C)** Volcano plots of DEGs in HCoV-OC43–, SARS-CoV-2–, and MERS-CoV–infected samples compared with uninfected control. Significantly upregulated genes (adjusted p-value < 0.05 and $\log_2$(fold change) > 1) are shown in red, while significantly downregulated genes (adjusted p-value < 0.05 and $\log_2$(fold change) < -1) are shown in blue. Antiviral target genes identified through our computational framework for each dataset are shown in black with annotation. For clear visualization, selected genes related to complex I are annotated. **(D)** Dot plots of GO enrichment results for significantly upregulated (p-value < 0.05 and $\log_2$(fold change) > 1) and downregulated (p-value < 0.05 and $\log_2$(fold change) < -1) DEGs. The top 10 biological processes are listed in descending order of adjusted p-value. Target subsystems (GO:0006119, oxidative phosphorylation; GO:0072529, pyrimidine-containing compound catabolic process) are highlighted in bold. **(E)** Rank plots of absolute NESs for Reactome pathways identified by GSEA (p-value cutoff = 0.99). Gray dots indicate the FDR for each pathway. Target subsystems (Complex IV assembly, R-HAS-9864848; Pyrimidine catabolism, R-HAS-73621) are indicated. Note that Complex I biogenesis (R-HSA-6799198) was not detected in the GSEA results for the HCoV-OC43–infected sample.

| GSE155986 (HCoV-229E) | | | |
|---|---|---|---|
| **Gene ID** | **Gene symbol** | **Gene name** | **Subsystem** |
| ENSG00000080819 | CPOX | coproporphyrinogen oxidase | Porphyrin metabolism |
| ENSG00000188690 | UROS | uroporphyrinogen III synthase | Porphyrin metabolism |
| ENSG00000256269 | HMBS | hydroxymethylbilane synthase | Porphyrin metabolism |
| ENSG00000126088 | UROD | uroporphyrinogen decarboxylase | Miscellaneous & Porphyrin metabolism |
| ENSG00000144659 | SLC25A38 | solute carrier family 25 member 38 | Transport reactions |
| ENSG00000143224 | PPOX | protoporphyrinogen oxidase | Heme synthesis & Porphyrin metabolism |
| ENSG00000148218 | ALAD | aminolevulinate dehydratase | Porphyrin metabolism |

**Table S1. List of predicted antiviral target genes in MRC-5 cells infected with HCoV-229E**

| GSE193169 (HCoV-OC43) | | | |
|---|---|---|---|
| **Gene ID** | **Gene symbol** | **Gene name** | **Subsystem** |
| ENSG00000169239 | CA5B | Carbonic anhydrase 5B | Transport reactions & Arginine and proline metabolism & Miscellaneous |
| ENSG00000131174 | COX7B | Cytochrome c oxidase subunit 7B | Oxidative phosphorylation |
| ENSG00000161281 | COX7A1 | Cytochrome c oxidase subunit 7A1 | Oxidative phosphorylation |
| ENSG00000135940 | COX5B | Cytochrome c oxidase subunit 5B | Oxidative phosphorylation |
| ENSG00000131055 | COX4I2 | Cytochrome c oxidase subunit 4I2 | Oxidative phosphorylation |
| ENSG00000131143 | COX4I1 | Cytochrome c oxidase subunit 4I1 | Oxidative phosphorylation |
| ENSG00000156885 | COX6A2 | Cytochrome c oxidase subunit 6A2 | Oxidative phosphorylation |
| ENSG00000170516 | COX7B2 | Cytochrome c oxidase subunit 7B2 | Oxidative phosphorylation |
| ENSG00000198804 | MT-CO1 | Mitochondrially encoded cytochrome c oxidase I | Oxidative phosphorylation |
| ENSG00000111775 | COX6A1 | Cytochrome c oxidase subunit 6A1 | Oxidative phosphorylation |
| ENSG00000198938 | MT-CO3 | Mitochondrially encoded cytochrome c oxidase III | Oxidative phosphorylation |
| ENSG00000176340 | COX8A | Cytochrome c oxidase subunit 8A | Oxidative phosphorylation |
| ENSG00000198712 | MT-CO2 | Mitochondrially encoded cytochrome c oxidase II | Oxidative phosphorylation |
| ENSG00000178741 | COX5A | Cytochrome c oxidase subunit 5A | Oxidative phosphorylation |
| ENSG00000160471 | COX6B2 | Cytochrome c oxidase subunit 6B2 | Oxidative phosphorylation |
| ENSG00000187581 | COX8C | Cytochrome c oxidase subunit 8C | Oxidative phosphorylation |
| ENSG00000127184 | COX7C | Cytochrome c oxidase subunit 7C | Oxidative phosphorylation |
| ENSG00000112695 | COX7A2 | Cytochrome c oxidase subunit 7A2 | Oxidative phosphorylation |
| ENSG00000164919 | COX6C | Cytochrome c oxidase subunit 6C | Oxidative phosphorylation |
| ENSG00000126267 | COX6B1 | Cytochrome c oxidase subunit 6B1 | Oxidative phosphorylation |

**Table S2. List of predicted antiviral target genes in A549-ACE2 cells infected with HCoV-OC43**

| | | GSE262439 (HCoV-OC43) | |
|---|---|---|---|
| Gene ID | Gene symbol | Gene name | Subsystem |
| ENSG00000198786 | MT-ND5 | mitochondrially encoded NADH:ubiquinone oxidoreductase core subunit 5 | Oxidative phosphorylation |
| ENSG00000109390 | NDUFC1 | NADH:ubiquinone oxidoreductase subunit C1 | Oxidative phosphorylation |
| ENSG00000115286 | NDUFS7 | NADH:ubiquinone oxidoreductase core subunit S7 | Oxidative phosphorylation |
| ENSG00000168653 | NDUFS5 | NADH:ubiquinone oxidoreductase subunit S5 | Oxidative phosphorylation |
| ENSG00000167792 | NDUFV1 | NADH:ubiquinone oxidoreductase core subunit V1 | Oxidative phosphorylation |
| ENSG00000160194 | NDUFV3 | NADH:ubiquinone oxidoreductase subunit V3 | Oxidative phosphorylation |
| ENSG00000170906 | NDUFA3 | NADH:ubiquinone oxidoreductase subunit A3 | Oxidative phosphorylation |
| ENSG00000198840 | MT-ND3 | mitochondrially encoded NADH:ubiquinone oxidoreductase core subunit 3 | Oxidative phosphorylation |
| ENSG00000198886 | MT-ND4 | mitochondrially encoded NADH:ubiquinone oxidoreductase core subunit 4 | Oxidative phosphorylation |
| ENSG00000213619 | NDUFS3 | NADH:ubiquinone oxidoreductase core subunit S3 | Oxidative phosphorylation |
| ENSG00000189043 | NDUFA4 | NDUFA4 mitochondrial complex associated | Oxidative phosphorylation |
| ENSG00000186010 | NDUFA13 | NADH:ubiquinone oxidoreductase subunit A13 | Oxidative phosphorylation |
| ENSG00000147684 | NDUFB9 | NADH:ubiquinone oxidoreductase subunit B9 | Oxidative phosphorylation |
| ENSG00000184752 | NDUFA12 | NADH:ubiquinone oxidoreductase subunit A12 | Oxidative phosphorylation |
| ENSG00000178127 | NDUFV2 | NADH:ubiquinone oxidoreductase core subunit V2 | Oxidative phosphorylation |
| ENSG00000183648 | NDUFB1 | NADH:ubiquinone oxidoreductase subunit B1 | Oxidative phosphorylation |
| ENSG00000174886 | NDUFA11 | NADH:ubiquinone oxidoreductase subunit A11 | Oxidative phosphorylation |
| ENSG00000198888 | MT-ND1 | mitochondrially encoded NADH:ubiquinone oxidoreductase core subunit 1 | Oxidative phosphorylation |
| ENSG00000212907 | MT-ND4L | mitochondrially encoded NADH:ubiquinone oxidoreductase core subunit 4L | Oxidative phosphorylation |
| ENSG00000136521 | NDUFB5 | NADH:ubiquinone oxidoreductase subunit B5 | Oxidative phosphorylation |
| ENSG00000090266 | NDUFB2 | NADH:ubiquinone oxidoreductase subunit B2 | Oxidative phosphorylation |
| ENSG00000165264 | NDUFB6 | NADH:ubiquinone oxidoreductase subunit B6 | Oxidative phosphorylation |
| ENSG00000147123 | NDUFB11 | NADH:ubiquinone oxidoreductase subunit B11 | Oxidative phosphorylation |
| ENSG00000131495 | NDUFA2 | NADH:ubiquinone oxidoreductase subunit A2 | Oxidative phosphorylation |
| ENSG00000119013 | NDUFB3 | NADH:ubiquinone oxidoreductase subunit B3 | Oxidative phosphorylation |
| ENSG00000110717 | NDUFS8 | NADH:ubiquinone oxidoreductase core subunit S8 | Oxidative phosphorylation |
| ENSG00000198695 | MT-ND6 | mitochondrially encoded NADH:ubiquinone oxidoreductase core subunit 6 | Oxidative phosphorylation |
| ENSG00000164258 | NDUFS4 | NADH:ubiquinone oxidoreductase subunit S4 | Oxidative phosphorylation |
| ENSG00000119421 | NDUFA8 | NADH:ubiquinone oxidoreductase subunit A8 | Oxidative phosphorylation |
| ENSG00000151366 | NDUFC2 | NADH:ubiquinone oxidoreductase subunit C2 | Oxidative phosphorylation |
| ENSG00000198763 | MT-ND2 | mitochondrially encoded NADH:ubiquinone oxidoreductase core subunit 2 | Oxidative phosphorylation |
| ENSG00000145494 | NDUFS6 | NADH:ubiquinone oxidoreductase subunit S6 | Oxidative phosphorylation |
| ENSG00000004779 | NDUFAB1 | NADH:ubiquinone oxidoreductase subunit AB1 | Oxidative phosphorylation |
| ENSG00000065518 | NDUFB4 | NADH:ubiquinone oxidoreductase subunit B4 | Oxidative phosphorylation |
| ENSG00000128609 | NDUFA5 | NADH:ubiquinone oxidoreductase subunit A5 | Oxidative phosphorylation |
| ENSG00000139180 | NDUFA9 | NADH:ubiquinone oxidoreductase subunit A9 | Oxidative phosphorylation |
| ENSG00000023228 | NDUFS1 | NADH:ubiquinone oxidoreductase core subunit S1 | Oxidative phosphorylation |
| ENSG00000130414 | NDUFA10 | NADH:ubiquinone oxidoreductase subunit A10 | Oxidative phosphorylation |
| ENSG00000140990 | NDUFB10 | NADH:ubiquinone oxidoreductase subunit B10 | Oxidative phosphorylation |
| ENSG00000125356 | NDUFA1 | NADH:ubiquinone oxidoreductase subunit A1 | Oxidative phosphorylation |
| ENSG00000099795 | NDUFB7 | NADH:ubiquinone oxidoreductase subunit B7 | Oxidative phosphorylation |
| ENSG00000158864 | NDUFS2 | NADH:ubiquinone oxidoreductase core subunit S2 | Oxidative phosphorylation |
| ENSG00000166136 | NDUFB8 | NADH:ubiquinone oxidoreductase subunit B8 | Oxidative phosphorylation |
| ENSG00000184983 | NDUFA6 | NADH:ubiquinone oxidoreductase subunit A6 | Oxidative phosphorylation |

### Table S3. List of predicted antiviral target genes in HBE organoids infected with HCoV-OC43

| GSE262439 (SARS-CoV-2) | | | |
| --- | --- | --- | --- |
| **Gene ID** | **Gene symbol** | **Gene name** | **Subsystem** |
| ENSG00000188641 | DPYD | dihydropyrimidine dehydrogenase | Pyrimidine metabolism |
| ENSG00000092964 | DPYSL2 | dihydropyrimidinase like 2 | Pyrimidine metabolism |
| ENSG00000100024 | UPB1 | beta-ureidopropionase 1 | Beta-alanine metabolism |
| ENSG00000101438 | SLC32A1 | solute carrier family 32 member 1 | Transport reactions |
| ENSG00000110090 | CPT1A | carnitine palmitoyltransferase 1A | Carnitine shuttle (cytosolic) & Carnitine shuttle (endoplasmic reticular) & Transport reactions & Carnitine shuttle (peroxisomal) & Fatty acid oxidation |

**Table S4. List of predicted antiviral target genes in HBE organoids infected with SARS-CoV-2**

| GSE262439 (MERS-CoV) | | | |
|---|---|---|---|
| **Gene ID** | **Gene symbol** | **Gene name** | **Subsystem** |
| ENSG00000188641 | DPYD | dihydropyrimidine dehydrogenase | Pyrimidine metabolism |
| ENSG00000092964 | DPYSL2 | dihydropyrimidinase like 2 | Pyrimidine metabolism |
| ENSG00000100024 | UPB1 | beta-ureidopropionase 1 | Beta-alanine metabolism |
| ENSG00000101438 | SLC32A1 | solute carrier family 32 member 1 | Transport reactions |

**Table S5. List of predicted antiviral target genes in HBE organoids infected with MERS-CoV**

| Series | Uninfected samples | Virus-infected samples | Remarks |
|---|---|---|---|
| GSE155986 | GSM4718629 ~ GSM4718631 | GSM4718635 ~ GSM4718637 | |
| GSE193169 | GSM6241725 ~ GSM6241727 | GSM6241731 ~ GSM6241733 | Uninfected = A549-ACE2 + sgCTRL |
| GSE262439 | GSM8169722 ~ GSM8169724 | GSM8169728 ~ GSM8169730<br>GSM8169731 ~ GSM8169733<br>GSM8169734 ~ GSM8169736 | Organoid scRNA-seq<br>(from our previous study) |

**Table S6. List of RNA-seq datasets used in this study**

| Gene | | Sequences (5' - 3') |
| --- | --- | --- |
| shSLC25A38 | Forward | CCGGGCTTATGTTACATCCGGTGATCTCGAGATCACCGGATGTAACATAAGCTTTTTG |
| | Reverse | AATTCAAAAAGCTTATGTTACATCCGGTGATCTCGAGATCACCGGATGTAACATAAGC |
| shALAD | Forward | CCGGGATGAGCTACAGTGCCAAATTCTCGAGAATTTGGCACTGTAGCTCATCTTTTTG |
| | Reverse | AATTCAAAAAGATGAGCTACAGTGCCAAATTCTCGAGAATTTGGCACTGTAGCTCATC |

**Table S7. List of shRNA oligomers**

| Gene | | Sequences (5' - 3') |
|---|---|---|
| ACTB | Forward | TCT GGC TCC TAG CAC CCT GA |
| | Reverse | CCA CCG ATC CAC ACA GAG TAC T |
| SLC25A38 | Forward | ACA CGG TGG AAA CGC TTA TGT TA |
| | Reverse | CCA GTG AGG CCA GAA TAC CA |
| ALAD | Forward | ACT TGG CAA CAG GGT ATC GG |
| | Reverse | AGC TGG GCT TGA CTT AGC TG |
| HCoV-OC43_N | Forward | AGC AAC CAG GCT GAT GTC AAT ACC |
| | Reverse | AGC AGA CCT TCC TGA GCC TTC AAT |
| HCoV-OC43_ORF1 | Forward | CAC TGT TGC TGG TGT TTC CA |
| | Reverse | GCG GCG TAA CAT ATC ATC CC |
| HCoV-229E_M | Forward | TTC CGA CGT GCT CGA ACT TT |
| | Reverse | CCA ACA CGG TTG TGA CAG TGA |
| HCoV-229E_ORF1 | Forward | TGG CAA ACA GCG TAT AAC CA |
| | Reverse | GAT CCT CGC TCC AGG GTA AT |
| SARS-CoV-2_N | Forward | GGC CAG AAG CTG GAC TTC CC |
| | Reverse | AGG ATT GCG GGT GCC AAT GT |
| SARS-CoV-2_ORF1 | Forward | GAT GAT GAT TAT TTC AAT AAA AAG GAC TGG TAT |
| | Reverse | GCG TGG TTT GTA TGA AAT CAC CG |
| MERS-CoV_N | Forward | GAC CCG AAG CAG CAC TCC CA |
| | Reverse | AGG GTT CCG CGT CCC AAA AGT T |
| MERS-CoV_ORF1 | Forward | AAT GCT GGG TGG AAA CCG AT |
| | Reverse | CAA AGC AAC CGG CAG ACA AA |

**Table S8. List of RT-qPCR primer sequences**
